# Beyond tracking climate: niche shifts during native range expansion and their implications for novel invasions

**DOI:** 10.1101/2021.06.09.447486

**Authors:** Nicky Lustenhouwer, Ingrid M. Parker

## Abstract

**Aim:** Although ecological niche models have been instrumental in understanding the widespread species distribution shifts under global change, rapid niche shifts limit model transferability to novel locations or time periods. Niche shifts during range expansion have been studied extensively in invasive species, but may also occur in native populations tracking climate change. We compared niche shifts during both types of range expansion in a Mediterranean annual plant to ask (i) whether the species’ native range expansion tracked climate change, (ii) whether further range expansion was promoted by niche expansion, and (iii) how these results changed forecasts of two ongoing invasions in Australia and California.

**Location:** Eurasian Holarctic, California and Australia

**Taxon:** *Dittrichia graveolens* (L.) Greuter (Asteraceae)

**Methods:** Niche shifts were quantified in both environmental and geographic space, using the framework of niche centroid shift, overlap, unfilling, and expansion (COUE) as well as Maximum Entropy modelling. We used the historic native distribution and climate data (1901-1930) to project the expected distribution in the present climate (1990-2019), and compared it to the observed current distribution of *D. graveolens*. Finally, we compared invasion forecasts based on the historic and present native niches.

**Results:** We found that *D. graveolens* expanded its native range well beyond what would be sufficient to track climate change, associated with a 5.5% niche expansion towards more temperate climates. In contrast, both invasions showed niche conservatism, and were (still) constrained to climatic areas matching the historic native niche.

**Main conclusions:** our results show that contrary to hypotheses in the literature, niche shifts are not necessarily more rapid in invasions than in native range expansions. We conclude that niche expansion during the process of climate tracking may cause further range expansion than expected based on climate change alone.

## Introduction

Forecasting the widespread distribution shifts of both native and invasive species under global change represents one of the major challenges in biodiversity conservation (Urban et al., 2016). The climate niche has become a central concept in modelling efforts to understand species’ preferred climate conditions, where such suitable habitat will be present under future climates, and which areas are at risk of invasion on other continents (Elith & Leathwick, 2009). Ecological niche models (ENMs; also known as habitat suitability models or species distribution models) are widely used to predict species distributions for scientific research and conservation planning (Araújo et al., 2011). The accuracy of these forecasts relies on the fundamental assumption of ENMs that niches are conserved in space and time (Pearman et al., 2008). However, niche shifts may occur for a variety of reasons, for example when species distributions are constrained by non-climatic factors (Alexander & Edwards, 2010). In addition, a growing body of empirical studies has demonstrated that range-expanding populations can rapidly adapt to novel environments (reviewed in Chuang & Peterson, 2016; Colautti & Lau, 2015), leading to evolutionary changes in the fundamental niche. Newly developed mechanistic and hybrid species distribution models that incorporate evolution of species’ physiological limits (Catullo et al., 2015; Hoffmann & Sgrò, 2011) predict markedly different outcomes of climate-induced range shifts (Bush et al., 2016) and invasions (Kearney et al., 2009) than traditional correlative ENMs. While there is thus a compelling argument for incorporating niche shifts into range expansion forecasts (Nadeau & Urban, 2019), a better understanding of the degree to which climate niche shifts promote contemporary range expansions is necessary to make informed predictions.

Niche shifts have been studied extensively in invasive species, with studies comparing the climate conditions occupied by populations in the native and invaded range. There is strong evidence that invading populations can rapidly evolve to re-establish adaptive clines along environmental gradients similar to those in their native range (e.g., Boheemen et al., 2019). Yet whether species’ ultimate niche limits are generally conserved during invasion (Liu et al., 2020; Petitpierre et al., 2012) or commonly shift (Atwater et al., 2018; Early & Sax, 2014) is highly debated. Niche stasis (*sensu* Pearman et al., 2008) is defined as the lack of change in either the fundamental niche (the climate where a species can grow in the absence of biotic constraints and geographic barriers) or realized niche (the actual climate conditions where a species is observed, which are captured by ENMs). Realized niches may shift in the invaded range when not all areas with similar climates are occupied (yet) due to dispersal limitation. Niche expansion occurs when biotic interactions or dispersal barriers constraining the realized niche in the native range are lifted in the new range, or when the fundamental niche itself evolves (Alexander & Edwards, 2010). Examples of evolutionary changes that have been linked to climate niche expansion in invasive populations include plant phenology responses to temperature or photoperiod (Colautti & Barrett, 2013; Dlugosch & Parker, 2008) and insect thermal and moisture tolerance (Hill et al., 2013; Kearney et al., 2009).

Much less attention in the empirical literature has been paid to niche shifts during contemporary native range shifts induced by climate change. Instead, species’ responses to global warming are commonly viewed as a “move, adapt, or die” conundrum, where populations need to track suitable climates to higher latitudes and altitudes, or adapt *in situ*, or else they will lag behind the pace of climate change and go extinct (Aitken et al., 2008; Corlett & Westcott, 2013). However, poleward-spreading populations face a variety of novel abiotic (as well as biotic) conditions, even if range expansion is initiated by climate change (Spence & Tingley, 2020). For example, photoperiod and temperature seasonality increase non-linearly with latitude, and plants experience a reduction in photosynthetically active radiation and light quality towards the poles (Saikkonen et al., 2012; Taulavuori et al., 2010). These novel combinations of temperature and photoperiod cues at higher latitudes require a plastic or evolutionary response (Visser, 2008). Thus, whether climate-mediated range shifts will involve simultaneous niche shifts is now acknowledged as an important open question (Lee-Yaw et al., 2019; Nadeau & Urban, 2019). Recent empirical examples of rapid evolutionary responses to novel environments during native range expansions include increased thermal niche breadth in damselflies (Lancaster et al., 2015) and earlier fall phenology in plants (Lustenhouwer et al., 2018).

Authors have hypothesized that niche shifts are more common or rapid in exotic than in native range expansions (Pearman et al., 2008; Wiens et al., 2019), because the realized niche in the native range would be more strongly limited by biotic interactions with predators, pathogens or competitors, whereas species would be able to occupy more of their fundamental niche in the introduced range where these biotic interactions are absent (Wiens et al., 2019). Strong genetic bottlenecks upon introduction combined with barriers to gene flow from the historic native range could also lead to high niche divergence in invading populations (Jakob et al., 2010). In the native range, the impact of ongoing gene flow on niche shifts will depend on whether that gene flow has a maladaptive swamping effect or rather increases evolutionary potential, an issue which is highly debated (Kottler et al., 2021). Comparing invasions to native range shifts can provide valuable insights into the drivers of niche shifts during range expansion in both native and exotic ranges. For example, quantifying niche expansion during invasions can help predict the degree of niche shifts that may be expected during native range expansions induced by climate change on similar time scales (Moran & Alexander, 2014; Wiens et al., 2019). Similarly, knowing whether a species’ realized niche in the native range is limited by biotic interactions and dispersal barriers or by fundamental evolutionary constraints such as genetic correlations will inform the likelihood of niche shifts in the invaded range (Alexander & Edwards, 2010).

In this study, we examine climate niche shifts during expansion of both native and exotic ranges by taking advantage of a species currently involved in both types of population spread. *Dittrichia graveolens* (L.) Greuter is an annual plant in the Asteraceae with a native distribution around the Mediterranean Basin (Brullo & De Marco, 2000). The species has greatly expanded its native range northward since the mid-20^th^ century, now occurring as far as Poland (Kocián, 2015). *D. graveolens* has invaded worldwide in most other regions with a Mediterranean climate - first Australia (1860s; Parsons & Cuthbertson, 2001) and South Africa (GBIF.org, 2020), then California (1980s; Preston, 1997), and most recently Chile (Santilli et al., 2021). In Australia and California, the species is considered a noxious weed of high management concern due to a combination of rapid spread and toxicity to livestock, impacts on native plant communities, and human skin allergies (Brownsey et al., 2013b; Parsons & Cuthbertson, 2001). In addition to the opportunity to compare native and exotic range expansion within the same species, *D. graveolens* is uniquely well suited for the study of niche shifts during range expansion. First, the species has a ruderal life history and produces large numbers of wind-dispersed seeds, facilitating spread along roads where biotic interactions with other plant species play a minor role in its ecological success. Dispersal and biotic barriers to spread are therefore relatively minor, potentially making changes in the fundamental niche more apparent. Second, previous work found that *D. graveolens*’ native range expansion coincided with rapid evolution of earlier flowering time at the leading edge (Lustenhouwer et al., 2018). This evolutionary response to shorter growing seasons suggests that a climatic niche shift may have contributed to the rapid range expansion of *D. graveolens*.

Using ecological niche models, we ask: (a) Did *D. graveolens* simply track climate change during the native range expansion, or was range expansion promoted by a climate niche expansion? (b) Is there evidence of niche shifts in the invaded ranges? (c) How does taking into account niche expansion during the native range expansion change invasion predictions for California and Australia? To answer these questions, we applied the COUE scheme of niche centroid shift, overlap, unfilling and expansion (Guisan et al., 2014) to *D. graveolens*’ native range expansion with climate change (comparing the periods 1901-1930 and 1990-2019) and to the two exotic range expansions. This method quantifies niches in environmental space and is widely used to study niche dynamics of invasive species. In addition, to explore niche shifts during range expansion in geographic space, we fit species distribution models using maximum entropy modelling (Maxent), which was specifically designed for presence-only data (Phillips et al., 2006). Based on prior evidence for rapid evolution of phenology at the northern range edge (Lustenhouwer et al., 2018), we hypothesized that *D. graveolens*’ climate niche expanded during native range expansion in Eurasia. We expected greater niche filling and greater niche expansion in Australia than in California, due to *D. graveolens*’ much longer invasion history in the former region. Finally, we hypothesized that forecasting the two invaded distributions based on the newly expanded native range would increase our invasion risk assessment to a wider range of climates and geographic areas.

## Materials and Methods

### Data collection

#### Occurrence data

We compiled presence-only species occurrence data for *Dittrichia graveolens* (L.) Greuter and its nomenclatural synonyms *Inula graveolens* (L.) Desf., *Cupularia graveolens* (L.) Godr. & Gren., and *Erigeron graveolens* (L.). We used the Holarctic Floral Kingdom (Cox, 2001) as our study region (split between Eurasia/North Africa for the native range and North America for one of the invaded ranges), to take into account the broadest range of environments available to *D. graveolens* in its evolutionary history and to facilitate projection of our models to other continents (Merow et al., 2013). This study region also allows for comparison to other studies using the same spatial extent (Petitpierre et al., 2012). Our primary data source was the Global Biodiversity Information Facility (~75% of occurrences, GBIF.org, 2020), supplemented by country-level species occurrence databases, standard floras, articles in botanical journals, and herbarium records (data sources are listed in Appendix 1 and further discussed in Appendix S1, Table S1). All citizen science records from iNaturalist (iNaturalist, 2020) were verified manually. We used QGIS v3.8.2 (QGIS Development Team, 2019) to combine and convert all data to the WGS84 coordinate system with decimal degrees latitude and longitude. Spatial grids (UTM, MTB, etc.) were imported as cell centroids. To increase sampling density across the study region, we also georeferenced localities without spatial coordinates (such as towns and other geographic features) using GEOLocate (Rios, 2020). We removed duplicate records and those located at (0,0) degrees or country centroids. The final (expanded) native range dataset included 8157 occurrences. To study niche shifts in the exotic range of *D. graveolens*, we selected the invasions in Australia and California because they are both well-documented (Calflora, 2021; using data from GBIF.org, 2020).

#### Defining the historic native range limit

*D. graveolens*’ historic native distribution is centred around the Mediterranean Basin, extending eastward to the Middle East and NW-India, and northward into central France (Brullo & De Marco, 2000; Tutin et al., 1976). The first records of a large-scale northward range expansion of *D. graveolens* appear for the 1950s in northern France, initially associated with salt mines and followed by abundant spread along roads (Parent, 2011; Wagenitz, 1966). We chose 1901-1930 as the pre-expansion time window for our study, which matches the earliest available information about *D. graveolens*’ distribution in floras of France (Bonnier & Layens, 1909; Coste, 1903; Rouy, 1903) and the Balkan Peninsula (Hayek & Markgraf, 1931), as well as the earliest monthly climate data (see next section). We used the botanical literature, dated species occurrences, and online databases to define a hypothesized *historic native range limit* pre-expansion (detailed description in Appendix S2). We considered administrative regions where *D. graveolens* is reported as a native species to be part of the historic native range (e.g. Tutin et al., 1976; von Raab-Straube, 2021), refining country-level data using other data sources.

#### Environmental predictors

All modelling and statistical analyses were conducted in R v4.0.0 (R Core Team, 2020), using code adapted from Di Cola et al. (2017), Guisan et al. (2017), and Smith (2020a). Monthly temperature and precipitation data were compiled from the Climatic Research Unit (CRU TS4.04, Harris et al., 2020) for 1901-1930 (past) and 1990-2019 (present), and used to calculate all 19 WORLDCLIM variables (Fick & Hijmans, 2017) for each time period (dismo package, Hijmans et al., 2017). In addition, we calculated the average total number of frost days for September-December (hereafter “frost”) for the same data sets. We selected 6 predictors (Table 1) based on the biology of *D. graveolens*, the Mediterranean and temperate climates characteristic of the expanded native range, and criteria limiting multicollinearity among variables: a pairwise Pearson correlation of 0.75 or less and a Variance Inflation Factor (usdm package, Naimi et al., 2014) below 5 (as recommended in Guisan et al., 2017). *D. graveolens* is a late-season annual plant, establishing a rosette in late spring and fruiting in autumn (Brownsey et al., 2013b), with earlier phenology occurring at higher latitudes (Lustenhouwer et al., 2018). In early life stages, precipitation is key to the growth of a tap root (Brownsey et al., 2013a), whereas plants are sensitive to cold and especially frost later in the year (Parsons & Cuthbertson, 2001), when the end of the growing season constrains plant fitness (Lustenhouwer et al., 2018). We considered climate variables representing temperature, precipitation, and their variability. Collinearity was evaluated over the entire Eurasian Holarctic. Based on *D. graveolens*’ biology, we discarded annual temperature and precipitation in favour of frost days during the reproductive period and precipitation in the driest and warmest quarters. Temperature of these quarters was highly correlated with the other selected variables and excluded.

**Table 1.**
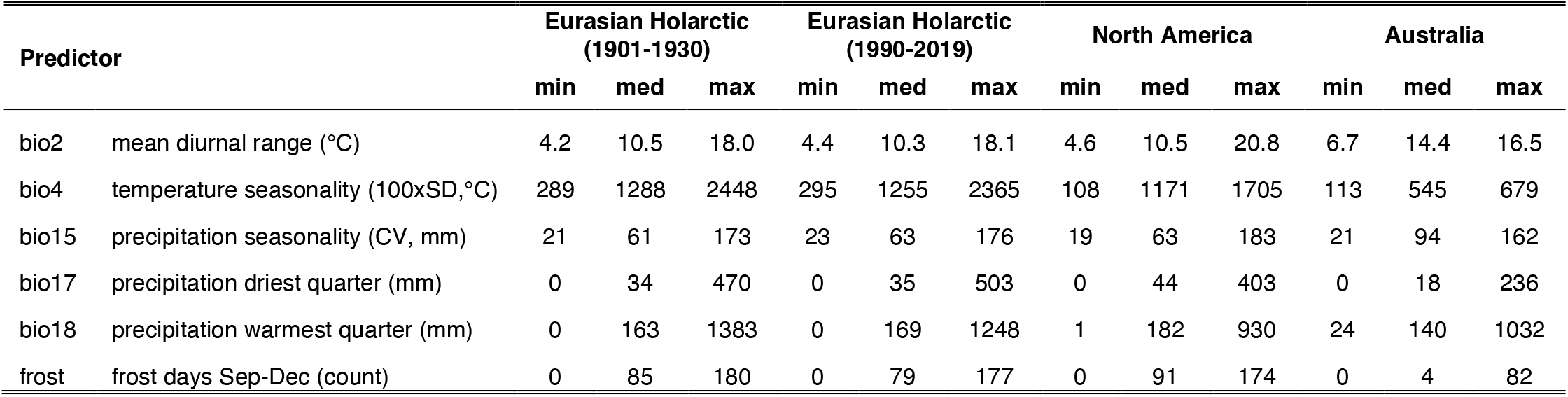
Environmental predictors included in this study, with minimum (min), median (med) and maximum (max) values across all pixels in each study area. Data from CRU TS4.04 (Harris et al., 2020).

| Predictor |  | Eurasian Holarctic<br>(1901-1930) |  |  | Eurasian Holarctic<br>(1990-2019) |  |  | North America |  |  | Australia |  |  |
| --- | --- | --- | --- | --- | --- | --- | --- | --- | --- | --- | --- | --- | --- |
|  |  | min | med | max | min | med | max | min | med | max | min | med | max |
| bio2 | mean diurnal range (°C) | 4.2 | 10.5 | 18.0 | 4.4 | 10.3 | 18.1 | 4.6 | 10.5 | 20.8 | 6.7 | 14.4 | 16.5 |
| bio4 | temperature seasonality (100xSD,°C) | 289 | 1288 | 2448 | 295 | 1255 | 2365 | 108 | 1171 | 1705 | 113 | 545 | 679 |
| bio15 | precipitation seasonality (CV, mm) | 21 | 61 | 173 | 23 | 63 | 176 | 19 | 63 | 183 | 21 | 94 | 162 |
| bio17 | precipitation driest quarter (mm) | 0 | 34 | 470 | 0 | 35 | 503 | 0 | 44 | 403 | 0 | 18 | 236 |
| bio18 | precipitation warmest quarter (mm) | 0 | 163 | 1383 | 0 | 169 | 1248 | 1 | 182 | 930 | 24 | 140 | 1032 |
| frost | frost days Sep-Dec (count) | 0 | 85 | 180 | 0 | 79 | 177 | 0 | 91 | 174 | 0 | 4 | 82 |

### Modelling approach

To combine climate and occurrence data, we scaled the latter down to the same resolution with one record per grid cell (0.5° latitude and longitude, corresponding to ca. 55 by 55 km). Cells containing occurrence records but no climate data (primarily covering sea rather than land, at coastlines or islands) were excluded. We set up two primary datasets, past and present, for our native range analyses covering the same Holarctic study region. The past dataset (representing the historic native range) consisted of climate data for the period 1901-1930 and all occurrence records located within the historic native range limit (n=399), assuming that occurrence locations represent suitable habitat for *D. graveolens* regardless of the date of observation. The present dataset (representing the expanded native range) consisted of climate data for the period 1990-2019 and the complete set of occurrence records (n=746). We used present-day climate data for the invaded range datasets. The Australia dataset contained all GBIF occurrences on the continent. The California dataset covered all of North America in spatial extent but occurrence data for California only, to focus on the west coast invasion.

#### Niches in environmental space

To quantify niche dynamics during range expansion following the COUE framework (Guisan et al., 2014), we applied the workflow developed by Broennimann et al. (2012), available in the ecospat R package (Broennimann et al., 2020; Di Cola et al., 2017). This approach evaluates niches in environmental space, which is defined by conducting a principal component analysis (PCA) for all environmental variables in both study regions under comparison. Niches are estimated by applying a kernel smoother function to the density of species occurrences from each range in gridded environmental space along the first two PCA axes. Indices of niche change are calculated exclusively for environments that are available in both study regions (using the 90^th^ percentile to exclude marginal environments), because shifts to and from nonanalog climates do not represent shifts in the fundamental niche (Guisan et al., 2014). *Niche stability* is defined as the proportion of occurrences in the new niche that overlap in environmental space with the native niche, while *niche expansion* (1-stability) is the proportion of occurrences in the new niche that are located in novel environments. Finally, *niche unfilling* quantifies environmental space that is occupied in the native range but that has not been filled in the new range (yet). It is calculated for the native occurrences as the proportion located in climate conditions that are unoccupied (but available) in the new range (Di Cola et al., 2017). Overall *niche overlap* is measured by Schoener’s *D* (Schoener, 1970) and used to test for niche conservatism during range expansion with two different tests (Broennimann et al., 2012; Warren et al., 2008). The niche equivalency test randomly reallocates occurrences between the two niches, creating a null distribution of *D* values based on 100 permutations to test whether the niches are identical (conducting one-sided tests asking if the observed *D* is greater or lower than expected by chance). The niche similarity test evaluates whether the two niches are more or less similar to each other than to other niches selected at random from the study area (shifting niches across both time periods for the native study area, but in the invaded study area only for Australia and California). We applied the above workflow to niche changes between (i) the past and present dataset (native range expansion), (ii) the past dataset and each invaded range, and (iii) the present dataset and each invaded range.

#### Projections in geographic space

To project niche changes in geographic space, we fit Maxent species distribution models (Phillips et al., 2006) to the past dataset (Past Model) and present dataset (Present Model). Maxent performs well for presence-only data under a range of sampling scenarios (Grimmett et al., 2020). Because the geographic availability of our native range occurrence data was highly concentrated in Western Europe, we used 3468 target background sites of taxonomically related species to correct for bias in sampling effort (Phillips et al., 2009; Appendix S3, Fig. S1). Models were fit using the maxnet package in R (Phillips, 2017). We first tuned the Past Model, using the trainMaxnet function (enmSdm package, A. B. Smith, 2020b) to select the optimal combination of features (starting with all classes) and regularization parameters (β of 0.5 and 1 to 10). The best model (lowest AIC) contained linear, quadratic and product features with β=0.5. We fit the Present Model using the same features and regularization. To ask whether *D. graveolens* simply tracked optimal climate conditions or expanded its native range beyond that, we created three model projections: the Past Model on the past and present climate, and the Present Model on the present climate. To evaluate model performance, we partitioned the past and present datasets randomly into training and test data using 5 k-folds. We computed AUC (dismo, Hijmans et al., 2017) and the Continuous Boyce Index (CBI; enmSdm, A. B. Smith, 2020b) for each k-fold and calculated the mean and standard deviation across models. To project areas at potential risk of invasion in California and Australia, we applied a Minimum Presence Threshold (the lowest habitat suitability at which *D. graveolens* is already present in the invaded range). We then compared projections generated by the Past and Present model for each invaded range.

## Results

Over the course of less than a century, *D. graveolens* has shifted its native range limit northward by nearly 7 degrees latitude. During this range expansion, the climate niche expanded by 5% to include more temperate environments with lower diurnal range and precipitation seasonality (bio2, bio15), increased precipitation in the driest and warmest quarters (bio17, bio18), and increased temperature seasonality (bio4) and fall frost (Fig. 1a,b). Niche overlap (Schoener’s *D* = 0.71) of the historic and expanded native niche was significantly lower than expected by chance (niche equivalency test; Fig. 1c, Table 2), indicating a niche shift during range expansion. Nonetheless, the two niches were more similar to each other than to randomly selected niches in the study area (niche similarity test; Fig. 1d, Table 2). Niche expansion was not driven by climate change between the past and present period, which happened in a different direction in environmental space (reduced frost and temperature seasonality; Fig. 1a).

**Figure 1.**
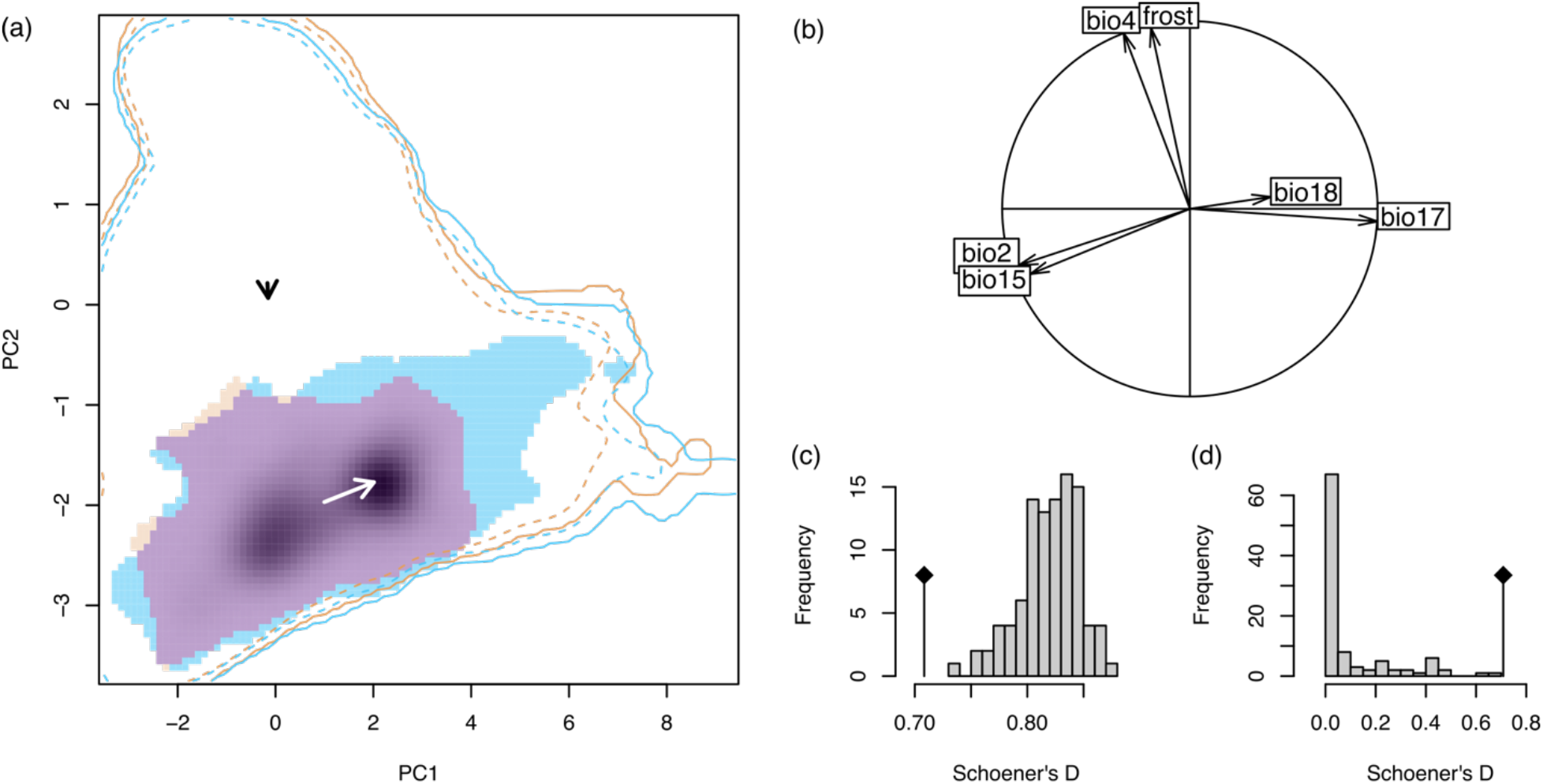
Niche shift during the native range expansion of *Dittrichia graveolens*, comparing the past (1901-1930) and present (1990-2019) datasets; (a) past and present niche in environmental space, with colours indicating niche stability (purple), unfilling (orange) and expansion (blue), and the white arrow showing the niche centroid shift. Dark shading shows the density of occurrences in the historic range. Lines represent the climate of the Eurasian Holarctic in the past (orange) and present (blue), using 100% (solid) and 90% (dashed) of available climates. The black arrow indicates the climate centroid shift between past and present. (b) principal component analysis of the entire environmental space, with axes explaining 41.6% (x) and 29.0% (y) of variation. Arrows show environmental predictors as defined in Table 1. (c, d) niche equivalency and niche similarity test, with histograms showing null distributions and diamonds marking observed niche overlap.

**Table 2.**
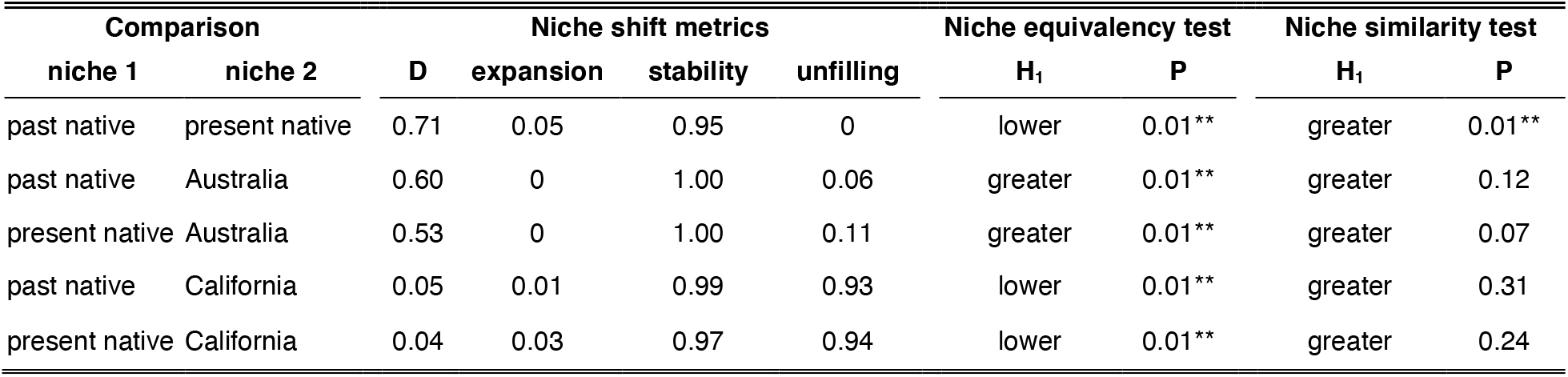
Niche shift metrics for *Dittrichia graveolens* following the COUE framework, calculated using the intersection of the 90^th^ percentile of environmental conditions in each range. Niche equivalency and similarity tests were one-sided, with H1 indicating the alternative hypothesis used.

| Comparison |  | Niche shift metrics |  |  |  | Niche equivalency test |  | Niche similarity test |  |
| --- | --- | --- | --- | --- | --- | --- | --- | --- | --- |
| niche 1 | niche 2 | D | expansion | stability | unfilling | H<sub>1</sub> | P | H<sub>1</sub> | P |
| past native | present native | 0.71 | 0.05 | 0.95 | 0 | lower | 0.01** | greater | 0.01** |
| past native | Australia | 0.60 | 0 | 1.00 | 0.06 | greater | 0.01** | greater | 0.12 |
| present native | Australia | 0.53 | 0 | 1.00 | 0.11 | greater | 0.01** | greater | 0.07 |
| past native | California | 0.05 | 0.01 | 0.99 | 0.93 | lower | 0.01** | greater | 0.31 |
| present native | California | 0.04 | 0.03 | 0.97 | 0.94 | lower | 0.01** | greater | 0.24 |

In line with this climate niche expansion, we found that *D. graveolens* in Eurasia has expanded its geographic range well beyond the extent sufficient to track climate change (Fig. 2, S2). The Past Model, fit to the historic climate and occurrences, predicts suitable habitat for *D. graveolens* around the Mediterranean Basin and into central France as expected (Fig. 2a; see Appendix S5, Fig. S3 for response functions for each predictor). Model AUC was 0.89 ± 0.01, indicating good to excellent performance (Araújo et al., 2005). Projecting this model onto the present climate (Fig. 2b), we found a minor northward shift in favourable conditions, particularly adjacent to the original northern range limit in France. However, the observed range expansion of *D. graveolens* extended much further northward and eastward (Fig. 2c). The Present Model, fit to the present climate and occurrences (Fig. S4), predicts a much wider distribution in Europe (Fig. 2d), outperforming the Past Model especially when predicting the actual probability of occurrence in the present (CBI of 0.93 and 0.61, respectively; Table 3).

**Figure 2.**
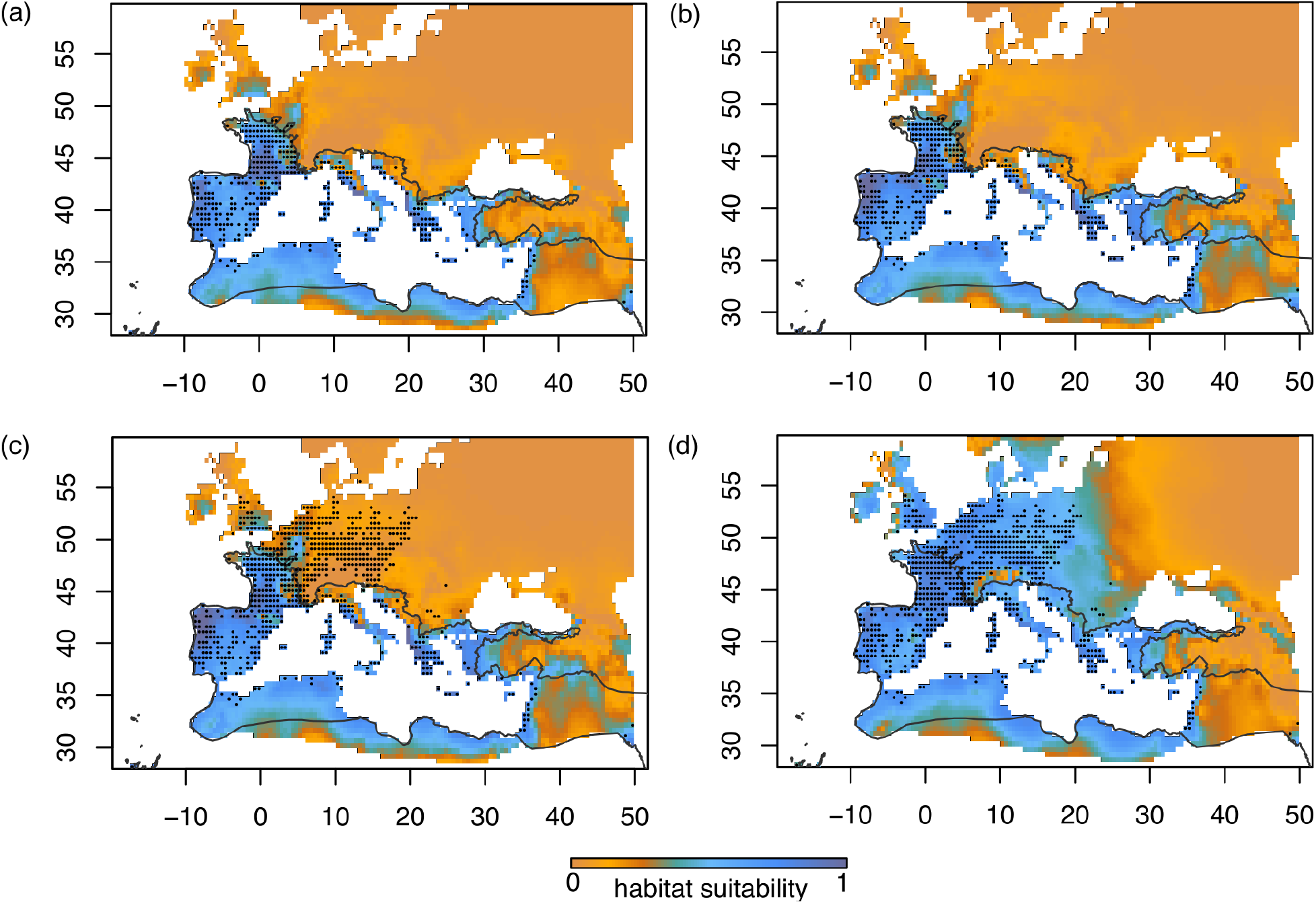
Maxent habitat suitability results for *Dittrichia graveolens*, using (a) the Past Model (fit to the past climate and historic native range occurrences) projected onto the past climate (1901-1930); (b) the Past Model projected onto the present climate (1990-2019), indicating expected range expansion with climate change; (c) the same projection with observed occurrences in the present, and (d) the Present Model (fit to the present climate and all occurrences) projected onto the present climate. Historic native range limit represented by the black line, and species occurrence records in the historic (a,b) and expanded (c,d) native range by dots. Axes display decimal degrees longitude (x) and latitude (y) using WGS84. Model projections onto the entire study area are available in Appendix S4, Fig. S2.

**Table 3.**
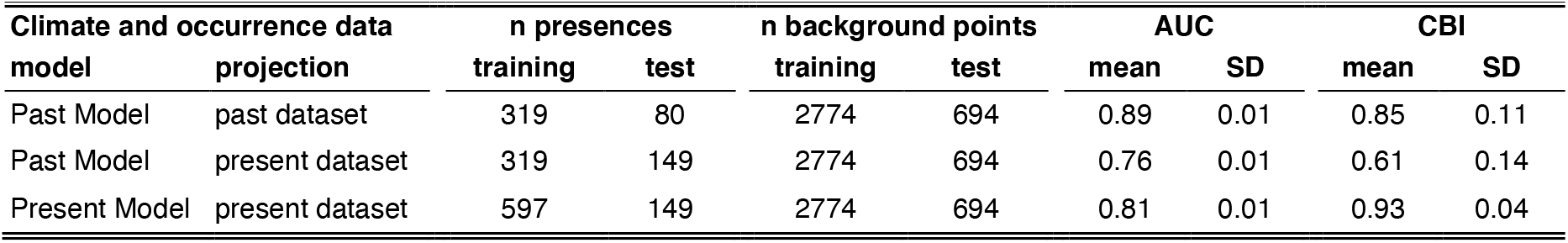
Maxent model performance for projections of the Past Model in both time periods, and the projection of the Present Model in the present. All presence and background points were allocated randomly to training and test datasets using 5 k-folds for cross-validation (table shows points per fold). Mean and standard deviation of AUC and CBI are given across the 5 k-folds.

| Climate and occurrence data |  | n presences |  | n background points |  | AUC |  | CBI |  |
| --- | --- | --- | --- | --- | --- | --- | --- | --- | --- |
| model | projection | training | test | training | test | mean | SD | mean | SD |
| Past Model | past dataset | 319 | 80 | 2774 | 694 | 0.89 | 0.01 | 0.85 | 0.11 |
| Past Model | present dataset | 319 | 149 | 2774 | 694 | 0.76 | 0.01 | 0.61 | 0.14 |
| Present Model | present dataset | 597 | 149 | 2774 | 694 | 0.81 | 0.01 | 0.93 | 0.04 |

The invasions in California and Australia exhibited contrasting niche dynamics, consistent with their difference in invader residence time. In California, only a small subset of the climate conditions in the native niche are already occupied (niche unfilling was 93% at the scale of North America; Fig. 3d, Table 2). In contrast, the Australian invasion has already spread into most areas that show similar climatic conditions to the native range (unfilling 6%; Fig. 3a, Table 2).

**Figure 3.**
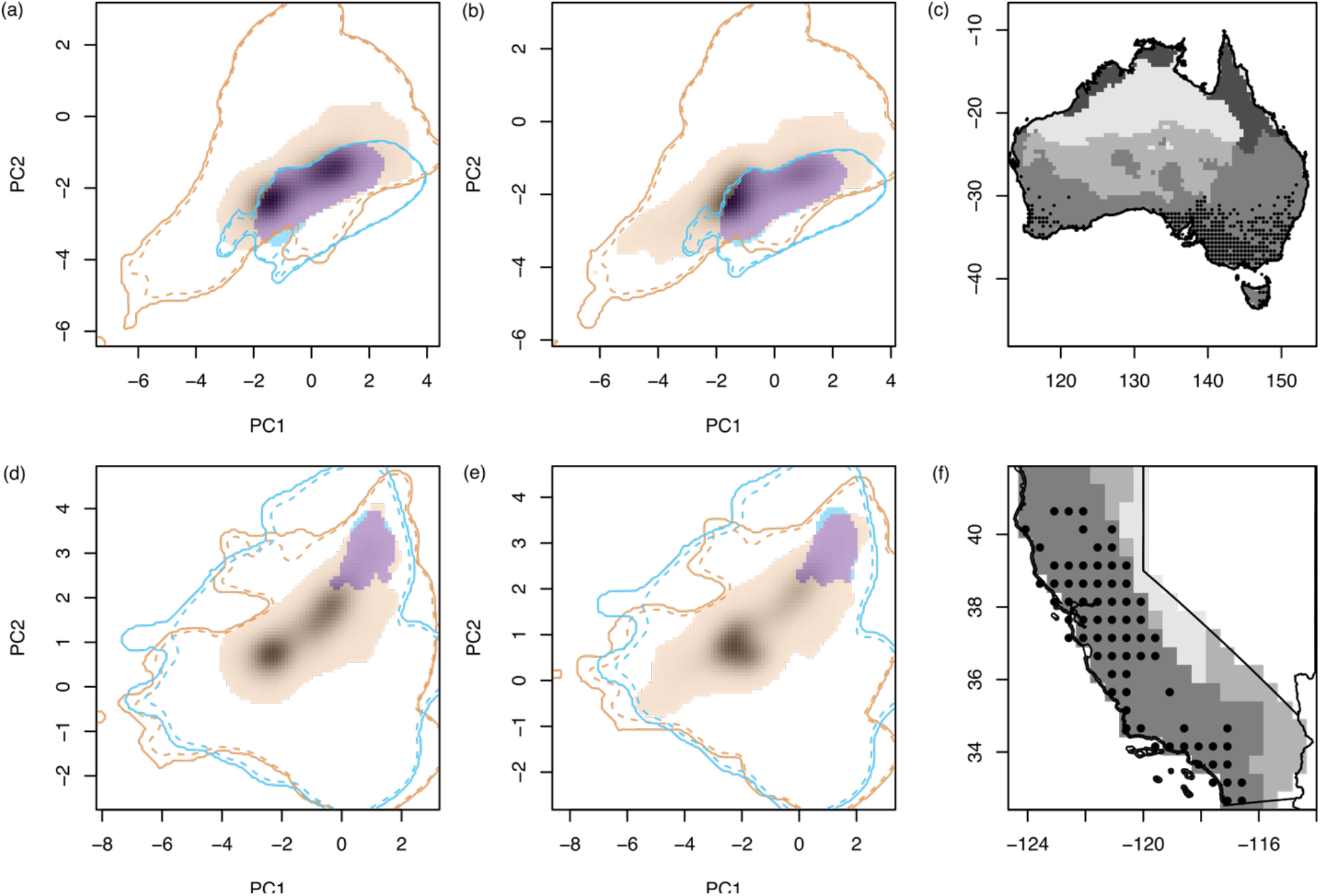
Niche shift during the invasion of *Dittrichia graveolens* in Australia (a,b) and California (d,e), using the past native (a,d) and present native (b,e) niche and climate as a reference. Colours indicate niche stability, unfilling and expansion as in Fig. 1, with lines representing the climate of the native (orange) and exotic (blue) study area. Shading shows the density of occurrences from the historic (a,d) and expanded (b,e) native range in environmental space. Panels c and f show areas at risk of invasion as projected by the Past and Present Model, with cells shaded from dark to light as follows: cell included in the Past Model only, in both models, in the Present Model only, or in neither model. Points indicate observation records for *D. graveolens* in Australia (c) and California (f), with axes displaying decimal degrees longitude (x) and latitude (y) using WGS84.

These contrasting results were reflected in the niche equivalency test, which showed that the California niche was less similar to the historic native range than expected by chance, while the Australian niche was more similar than expected (Table 2). We found near-complete niche stability in both invasions, meaning that invasive populations occur almost exclusively in climate conditions that also existed in the historic native range. Accordingly, niche expansion was low (Table 2). Niche conservatism could not be rejected in the niche similarity test for either invasion, indicating that it cannot be ruled out that any niche differentiation between native and exotic ranges was driven by the availability of environments in the exotic ranges.

Niche change indices were similar for both invasions when including the native range expansion (Table 2). Even though the peak of native occurrences shifted toward more temperate climates (shading in Fig. 3e), California niche unfilling remained stable because the majority of native occurrences already occurred outside the invaded climate space. Of the newly invadable climate conditions, most are absent in Australia (Fig. 3b), but present in North America (Fig. 3e). Combining projections of the Past and Present Maxent models onto Australia and California revealed additional areas that would be at risk of invasion if niche shifts were to happen as in the native range. However, neither invasion has yet advanced into areas solely included by the Present Model (Fig. 3c, 3f).

## Discussion

As many native species are shifting their ranges to track climate change, the ecological and evolutionary drivers of range expansion have become a key focus of biogeographical research (Nadeau & Urban, 2019). Ecological studies of native range shifts have found varying degrees to which species keep up with shifting climate isotherms (Chen et al., 2011; Lenoir et al., 2020), including some species with range shifts that outpace climate warming. However, few studies have explicitly studied whether a climatic niche shift occurred during contemporary native range shifts. Two studies using a similar approach to ours found incomplete climate tracking (*Ilex aquifolium*, Walther et al., 2005) or range contractions rather than expansions (montane rodents, Pardi et al., 2020). In a species that did expand its native range very rapidly, the butterfly *Acraea terpsicore*, a recent study found that new populations mostly occupied climatic areas similar to those in the historic native range, with little evidence for niche shifts (Chowdhury et al., 2021). In contrast, our results show that *D. graveolens* has undergone an extensive northward expansion of its native range, well beyond what would be expected based on climate tracking alone.

This native range expansion was accompanied by a 5.5% expansion of the climate niche (following Broennimann et al., 2012), a high number relative to studies of niche shifts during exotic range expansions using the same metrics for invasive species. Our results fall in the 78th percentile of niche expansion estimates by Petitpierre et al. (2012, n = 50 plant species), and the 60th percentile of species included by Liu et al. (2020, n = 211 plant species with COUE estimates). However, Early & Sax (2014) found much higher niche expansion overall than Petitpierre et al., which they attributed to life history differences between the study species; Petitpierre et al. studied weedy species that were widely distributed in their native range, similar to *D. graveolens*, while Early & Sax considered endemic species with small native ranges. The native ranges of these species may have been constrained by non-climatic factors, allowing for higher niche expansion in the introduced range. Understanding the drivers of realized niche shifts will be crucial to predicting both native and exotic range expansions (Early & Sax, 2014).

Expansion of the realized niche as quantified in this study can result from the removal of dispersal or biotic constraints, or be caused by evolution of the fundamental niche itself. The biology of *D. graveolens* and previous empirical work on its native range expansion provide insight into the possible role played by these drivers. Concerning dispersal limitation, *D. graveolens* has very high spread potential due to a combination of its annual life history, wind-dispersed seeds and the production of tens of thousands of seeds per plant (Lustenhouwer et al., 2018). No historic geographic barriers to dispersal are apparent in the native range. From a biotic perspective, *D. graveolens* has a typical ruderal life history and has expanded its native range primarily along roadsides, where species interactions are less predominant. Finally, theory predicts that rapid evolution of both locally adapted phenotypes and increased dispersal during range expansion could cause populations to spread beyond shifting climate isotherms, while expanding their fundamental niche to include colder climates (Kubisch et al., 2013). At the start of the native range expansion, the first records of *D. graveolens* north of the historic range limit in France follow the availability of new suitable habitat made available by climate change (Fig. 2b). Previous work on *D. graveolens* demonstrated that over the course of further range expansion to the Netherlands, rapid evolution of earlier phenology increased plant fitness in northern leading-edge populations, where reproduction is constrained by the earlier end of the growing season (Lustenhouwer et al., 2018). Adaptation to climate conditions at northern latitudes may therefore have contributed to the niche expansion found in this study (Fig. 1), facilitating the spread of *D. graveolens* beyond expectations based on climate tracking alone (Fig. 2c).

In contrast to the native range expansion, we found little evidence for niche expansion in the invaded range. Australia and California differed strongly in niche unfilling, illustrating the importance of invader residence time in invasion risk assessment (Wilson et al., 2007). Although the California invasion appears to be in much earlier stages with further spread expected, both invasive niches almost exclusively cover climatic conditions that are also present in the historical native niche (high stability), a pattern known as climate matching, which has been found for many successful invasive species (Hayes & Barry, 2008). Despite its longer history, the Australian niche barely expands beyond the native niche, suggesting niche conservatism. Our finding of greater niche expansion during native range expansion than during invasion contradicts a common hypothesis in the literature that niche stasis should be more pronounced in the native range (Pearman et al., 2008; Wiens et al., 2019). However, this hypothesis is based on the idea that biotic interactions are more limiting in the native range. It may be that biotic interactions had very little to do with *D. graveolens*’ range expansion with climate change, providing “an exception that proves the rule.”

Incorporating the native range expansion into projections of niche change and invasion risk adds newly invadable climatic areas, especially in North America (Fig. 3). However, given that neither invasion has yet spread into these areas, it remains an open question whether any similar niche expansion can be expected in the invaded ranges in the future. Population genetic analysis of samples collected widely across the native range would be needed to definitively identify the origin of the invasions in California and Australia. However, the strong overlap in climate conditions occupied in the historic native range and both exotic ranges does suggest that invasive populations originated from a Mediterranean part of the native range. Future studies should compare plants of native and invasive origin in order to assess the evolutionary potential of *D. graveolens* in California and Australia, and to predict whether these invasions will proceed into areas forecasted by the Present Model (Fig. 3c,f). Quantifying genetic variance in relevant traits and identifying genetically based changes from the origin to the leading edge within each invasion would be most informative of any ongoing and potential future niche shifts. Assuming that the evolution of locally adapted phenology clines has contributed to the expansion of *D. graveolens* in its native range (Lustenhouwer et al. 2018), the introduction of genotypes from northern Europe to the invaded ranges could put new areas at risk of invasion by this weed of concern. Our study therefore supports existing calls to limit multiple introductions of invasive species, even if they are widespread already (A. L. Smith et al., 2020).

### Limitations

Any niche modelling study is constrained by the available data. Although the native and invaded distributions of *D. graveolens* are relatively well-documented, occurrence records do contain temporal and spatial biases that may have affected our results. Because the majority of our occurrence data was reported in recent decades, we included all occurrences south of the original range limit in the historic native range, regardless of the date they were reported. It is possible that past climate conditions at these locations were less favourable for *D. graveolens*. However, because our study is focused on a northward range expansion tracking climate change, and the historic native range now represents the trailing edge of the distribution, our assumption to treat all occurrences as historical presences should underestimate the niche shift during range expansion and therefore represents a conservative approach. To address the spatial bias in our data, we employed the target background record approach to reflect sampling effort across the study area. This solution was a great improvement over randomly chosen background points (which resulted in overfitting to climate areas with high sampling effort), but was still not optimal given the low number of target background points available compared to Maxent’s standard of 10,000. However, at our spatial resolution (0.5°) even near 215,000 target records from 296 species covered only 3468 cells across the entire study area (Fig. S1). Higher-resolution climate data would allow for more background points (Merow et al., 2013) but was not available for the past time period of interest.

### Conclusion

Our results suggest that climate change may act as a catalyst for range expansion and subsequent climate niche expansion at higher latitudes in plants. The generality of this phenomenon for other species will depend on their range-limiting factors, phenotypic plasticity, evolutionary potential (generation time, heritable genetic diversity), and dispersal ability (Catullo et al., 2015), with short-living, ruderal plants similar to *D. graveolens* being likely candidates for niche expansion. Recent work on invertebrates suggests that range expansion can promote increased thermal and diet niche breadth at the range edge in a wide range of insect species (Lancaster, 2016, 2020; Neu et al., 2021). We encourage future studies validating climate-based projections of native range expansions with observed spread (e.g., Araújo et al., 2005), to examine whether climate niche shifts during native range expansion are common and cause populations to spread further than expected under climate tracking alone. Ultimately, the study of both biological invasions and native species threatened by climate change will benefit from a better understanding of the drivers of niche shifts during range expansion.

## Acknowledgements

We are grateful to Zarina Pringle for her assistance in the occurrence data collection. We also thank A. Pliszko, M. Kaligarič, G. Király, P. Eliáš, and D. Schmidt for pointing us to occurrence records in Poland, Slovenia, Slovakia, and Hungary. Finally, we thank the Botanical Society of Britain and Ireland for providing data from the UK. Members of the Parker and Gilbert labs provided helpful comments at several points in the study. This work was funded by Swiss National Science Foundation grant P2EZP3_178481 to N.L. and USDA NIFA grant 2020-67013-31856 to I.M.P.

## Data Availability

Data and R code needed to reproduce the results in the manuscript are available on the following GitHub repository: https://github.com/nickylustenhouwer/beyond-tracking-climate.

Further details on the occurrence data collection are provided in the Supporting Information (Appendix S1, Table S1).

## Biosketch

**Nicky Lustenhouwer** is an evolutionary ecologist studying how populations respond to rapidly changing environments, with a main focus on plant population spread. This work is part of her postdoctoral research at UC Santa Cruz studying evolution during the native and exotic range expansion of *Dittrichia graveolens*.

**Ingrid M. Parker** is a professor in plant evolutionary ecology at UC Santa Cruz. Her research interests include the invasion of non-native species, plant disease ecology, the evolution of domestication, ecological restoration, and plant conservation.

### Author contributions

N.L. and I.M.P. designed the study. N.L. collected and analysed the data and wrote the manuscript, with I.M.P. contributing revisions.

