## Supporting Information for "Beyond tracking climate: niche shifts during native range expansion and their implications for novel invasions"

**Contents**

| | |
|---|---|
| Appendix S1 | Species occurrence data collection |
| | Table S1 |
| Appendix S2 | Defining the historic native range limit |
| Appendix S3 | Target background point selection |
| | Figure S1 |
| Appendix S4 | Maxent model projections for the entire Eurasian Holarctic |
| | Figure S2 |
| Appendix S5 | Maxent response functions for each predictor |
| | Figure S3, S4 |

All citations are listed in the main manuscript under References or Appendix 1 (Data sources).

### Appendix S1: Species occurrence data collection

Species occurrence records for *Dittrichia graveolens* (L.) Greuter, used in this study are available in processed form at a resolution of 0.5° latitude/longitude on the following GitHub repository: https://github.com/nickylustenhouwer/beyond-tracking-climate.
These processed data were used as input for all analyses, which can be reproduced using the R scripts in the repository. Species occurrence records were compiled from the following sources:

*Global databases*

- Global Biodiversity Information Facility (GBIF.org, 2020)
- iNaturalist citizen science website, all records verified manually (iNaturalist, 2020)

*Country/state-level species occurrence databases and papers*

- Austria: Beitrage zur Flora von Österreich, IV (Stöhr et al., 2012)
- California: Calflora (Calflora, 2021)
- Croatia: Flora Croatica (Nikolić, 2015)
- Czech Republic: Distributions of vascular plants in the Czech Republic. Part 7 (Kaplan et al., 2018)
- Germany: FloraWeb (FloraWeb, 2013)
- Poland: ATPOL (Zając & Zając, 2019)
- Slovenia: map of *Dittrichia graveolens* as a new alien (Frajman & Kaligarič, 2009)
- Switzerland: Info Flora (Info Flora, 2020)
- United Kingdom: Botanical Society of Britain and Ireland database (BSBI, 2020)

*Individually processed records*

Our dataset contains additional records that are not included in the existing databases listed above. These records include manually verified iNaturalist observations not present in the GBIF dataset [1], and newly georeferenced GBIF records that contained locality descriptions but no spatial coordinates. We also compiled and georeferenced records from herbaria, standard floras, and botanical papers. All individually processes records are available in Table S1 below.

**Table S1** Individually processed species occurrence records for *Dittrichia graveolens* (L.) Greuter. Identifiers are gbifID, iNatID, or herbarium specimen ID when applicable.

| Longitude | Latitude | Country | Source type | Citation | Identifier * |
|---|---|---|---|---|---|
| 70.043241 | 34.285275 | Afghanistan | Flora | (Georgiadou et al., 1980) | |
| 69.760047 | 34.589615 | Afghanistan | Flora | (Georgiadou et al., 1980) | |
| 70.457365 | 34.428589 | Afghanistan | Flora | (Georgiadou et al., 1980) | |
| 70.764046 | 34.715809 | Afghanistan | Flora | (Georgiadou et al., 1980) | |
| 69.958581 | 36.763198 | Afghanistan | Flora | (Georgiadou et al., 1980) | |
| 69.21236 | 34.56585 | Afghanistan | Flora | (Georgiadou et al., 1980) | |
| 69.533559 | 36.734133 | Afghanistan | Flora | (Georgiadou et al., 1980) | |
| 69.949046 | 36.77955 | Afghanistan | Herbarium | (GBIF.org, 2020) | gbifID 665711331 |
| 69.533559 | 36.734133 | Afghanistan | Herbarium | (GBIF.org, 2020) | gbifID 665711332 |
| 69.947463 | 36.778277 | Afghanistan | Herbarium | (GBIF.org, 2020) | gbifID 665711331 |

| Longitude | Latitude | Country | Source type | Citation | Identifier * |
|---|---|---|---|---|---|
| 69.207486 | 34.555349 | Afghanistan | Herbarium | (Open Herbarium, 2020) | CoAF 009662 |
| 70.621679 | 34.171831 | Afghanistan | Herbarium | (Open Herbarium, 2020) | CoAF 014768 |
| 70.811995 | 36.734772 | Afghanistan | Herbarium | (Open Herbarium, 2020) | CoAF 000327 |
| 19.781016 | 41.777697 | Albania | Paper | (Shallari et al., 1998) | |
| -0.641667 | 35.691111 | Algeria | Herbarium | (GBIF.org, 2020) | gbifID 2517059524 |
| 0.089176 | 35.931151 | Algeria | Herbarium | (GBIF.org, 2020) | gbifID 2514123220 |
| -0.635936 | 35.698778 | Algeria | Herbarium | (GBIF.org, 2020) | gbifID 2435566178 |
| 3.087562 | 36.731875 | Algeria | Herbarium | (GBIF.org, 2020) | gbifID 1842855234 |
| 6.614722 | 36.365 | Algeria | Herbarium | (GBIF.org, 2020) | gbifID 1455444497 |
| -0.314678 | 35.847729 | Algeria | Herbarium | (GBIF.org, 2020) | gbifID 1096871738 |
| 6.797126 | 36.793988 | Algeria | Citizen science | (iNaturalist, 2020) | iNatID 47018949 |
| 6.692198 | 36.41339 | Algeria | Citizen science | (iNaturalist, 2020) | iNatID 35461843 |
| 11.45889 | 47.26583 | Austria | Paper | (Pagitz & Lechner-Pagitz, 2015) | |
| 11.39528 | 47.21611 | Austria | Paper | (Pagitz & Lechner-Pagitz, 2015) | |
| 11.16583 | 47.28417 | Austria | Paper | (Pagitz & Lechner-Pagitz, 2015) | |
| 10.93833 | 47.27944 | Austria | Paper | (Pagitz & Lechner-Pagitz, 2015) | |
| 11.19306 | 47.28167 | Austria | Paper | (Pagitz & Lechner-Pagitz, 2015) | |
| 11.24111 | 47.2625 | Austria | Paper | (Pagitz & Lechner-Pagitz, 2015) | |
| 3.449196 | 50.56765 | Belgium | Herbarium | (GBIF.org, 2020) | gbifID 1839780897 |
| 4.911539 | 50.468902 | Belgium | Herbarium | (GBIF.org, 2020) | gbifID 1455414896 |
| 18.18531 | 44.08668 | Bosnia-Herzegovina | Herbarium | (GBIF.org, 2020) | gbifID 1096873635 |
| 23.71671 | 42.77015 | Bulgaria | Paper | (Vladimirov et al., 2010, p. 14) | |
| 23.1144 | 42.8133 | Bulgaria | Paper | (Vladimirov et al., 2010, p. 14) | |
| 23.11969 | 42.58879 | Bulgaria | Paper | (Vladimirov & Tan, 2016, p. 31) | |
| 26.95872 | 42.97057 | Bulgaria | Paper | (Vladimirov et al., 2018, p. 37) | |
| 15.612732 | 43.797915 | Croatia | Paper | (Pandža, 1998) | |
| 9.473235 | 42.841692 | France | Citizen science | (iNaturalist, 2020) | iNatID 33418213 |
| 10.1963 | 52.11401 | Germany | Citizen science | (iNaturalist, 2020) | iNatID 33371865 |
| 12.929722 | 48.96611 | Germany | Citizen science | (iNaturalist, 2020) | iNatID 32215682 |
| 12.931577 | 48.970392 | Germany | Citizen science | (iNaturalist, 2020) | iNatID 32214764 |
| 12.481829 | 50.786573 | Germany | Citizen science | (iNaturalist, 2020) | iNatID 38025374 |
| 24.112649 | 41.09168 | Greece | Citizen science | (iNaturalist, 2020) | iNatID 16385019 |
| 16.6236088 | 47.2758016 | Hungary | Paper | (Schmidt, 2019) | |
| 7.712761 | 47.647838 | Hungary | Paper | (Takács et al., 2016) | |
| 17.2901511 | 47.6257761 | Hungary | Database | (OpenBioMaps, 2021) | |
| 76.517835 | 32.080729 | India | Herbarium | (GBIF.org, 2020) | gbifID 1563400212 |
| 54.443907 | 36.8427 | Iran | Flora | (Georgiadou et al., 1980) | |
| 49.84 | 32.0686111 | Iran | Flora | (Georgiadou et al., 1980) | |
| 48.67717 | 31.312381 | Iran | Flora | (Georgiadou et al., 1980) | |

| Longitude | Latitude | Country | Source type | Citation | Identifier * |
|---|---|---|---|---|---|
| 54.549038 | 28.755141 | Iran | Flora | (Georgiadou et al., 1980) | |
| 54.443907 | 36.8427 | Iran | Herbarium | (GBIF.org, 2020) | gbifID 1455977907 |
| 60.8966 | 29.572 | Iran | Herbarium | (GBIF.org, 2020) | gbifID 1935969854 |
| 57.30841 | 37.49302 | Iran | Paper | (Ghahremaninejad et al., 2012) | FUMH 21091 |
| 12.492252 | 41.89025 | Italy | Paper | (Caneva et al., 2002) | |
| 14.346637 | 37.025235 | Italy | Citizen science | (iNaturalist, 2020) | iNatID 8224589 |
| 16.067561 | 39.182501 | Italy | Herbarium | (De Stefani, 1973) | PAL 108059 |
| 11.25 | 43.766667 | Italy | Herbarium | (GBIF.org, 2020) | gbifID 2598858193 |
| 15.54351 | 38.188396 | Italy | Herbarium | (GBIF.org, 2020) | gbifID 2517552148 |
| 15.001 | 40.405 | Italy | Herbarium | (GBIF.org, 2020) | gbifID 2516939863 |
| 9.11315 | 39.240851 | Italy | Herbarium | (GBIF.org, 2020) | gbifID 2514965545 |
| 14.2167 | 42.4667 | Italy | Herbarium | (GBIF.org, 2020) | gbifID 2514559665 |
| 7.673054 | 45.076313 | Italy | Herbarium | (GBIF.org, 2020) | gbifID 2430805326 |
| 12.372159 | 43.109497 | Italy | Herbarium | (GBIF.org, 2020) | gbifID 2284179540 |
| 17.933333 | 40.633333 | Italy | Herbarium | (GBIF.org, 2020) | gbifID 1096873859 |
| 10.961295 | 44.424411 | Italy | Herbarium | (GBIF.org, 2020) | gbifID 1096873618 |
| 7.65 | 43.8 | Italy | Herbarium | (GBIF.org, 2020) | gbifID 1096873158 |
| 12.542384 | 38.014399 | Italy | Herbarium | (GBIF.org, 2020) | gbifID 1096872628 |
| 7.723705 | 43.809546 | Italy | Herbarium | (GBIF.org, 2020) | gbifID 1096872295 |
| 8.95 | 44.416667 | Italy | Herbarium | (GBIF.org, 2020) | gbifID 1096843040 |
| 8.333333 | 44.333333 | Italy | Herbarium | (GBIF.org, 2020) | gbifID 1096817020 |
| 13.35312 | 38.113415 | Italy | Herbarium | (GBIF.org, 2020) | gbifID 1096816988 |
| 14.070973 | 40.819235 | Italy | Herbarium | (GBIF.org, 2020) | gbifID 1096816978 |
| 14.262416 | 40.860949 | Italy | Herbarium | (GBIF.org, 2020) | gbifID 1096875492 |
| 12.483333 | 41.9 | Italy | Herbarium | (GBIF.org, 2020) | gbifID 1096873624 |
| 35.193889 | 33.273333 | Lebanon | Flora | (Mouterde, 1983) | |
| 35.375636 | 33.559932 | Lebanon | Flora | (Mouterde, 1983) | |
| 35.637778 | 33.978333 | Lebanon | Flora | (Mouterde, 1983) | |
| 35.510826 | 33.877621 | Lebanon | Flora | (Mouterde, 1983) | |
| 35.739931 | 34.35573 | Lebanon | Flora | (Mouterde, 1983) | |
| 35.849722 | 34.436667 | Lebanon | Flora | (Mouterde, 1983) | |
| 35.792778 | 33.993611 | Lebanon | Flora | (Mouterde, 1983) | |
| 35.904167 | 33.849722 | Lebanon | Flora | (Mouterde, 1983) | |
| 35.841667 | 33.936944 | Lebanon | Flora | (Mouterde, 1983) | |
| 35.785278 | 34.121944 | Lebanon | Flora | (Mouterde, 1983) | |
| 35.820833 | 33.795 | Lebanon | Flora | (Mouterde, 1983) | |
| 35.2038 | 33.2705 | Lebanon | Paper | (Tohmé & Tohmé, 2015) | |
| 35.3729 | 33.5571 | Lebanon | Paper | (Tohmé & Tohmé, 2015) | |
| 71.578488 | 34.008 | Pakistan | Flora | (Georgiadou et al., 1980) | |

| Longitude | Latitude | Country | Source type | Citation | Identifier * |
|---|---|---|---|---|---|
| 68.530907 | 30.241953 | Pakistan | Flora | (Open Herbarium, 2020) | FoPK 2101627 |
| 71.524915 | 34.015137 | Pakistan | Flora | (Open Herbarium, 2020) | FoPK 2101625 |
| 66.931671 | 24.940072 | Pakistan | Herbarium | (GBIF.org, 2020) | gbifID 1456108352 |
| 18.70894 | 49.76667 | Poland | Paper | (Kocián, 2015) | |
| 18.70992 | 49.76669 | Poland | Paper | (Kocián, 2015) | |
| 18.7105 | 49.76699 | Poland | Paper | (Kocián, 2015) | |
| 19.87788 | 49.91735 | Poland | Paper | (Pliszko & Kocián, 2017) | |
| 24.69377 | 45.79124 | Romania | Paper | (Szatmari & Hurdu, 2021) | |
| 24.60014 | 45.77224 | Romania | Paper | (Szatmari & Hurdu, 2021) | |
| 22.171269 | 43.301835 | Serbia | Paper | (Zlatković & Bogosavljević, 2014) | BEOU 16839 |
| 22.656946 | 43.025692 | Serbia | Paper | (Zlatković & Bogosavljević, 2014) | BEOU 16840 |
| 16.987324 | 48.679258 | Slovakia | Paper | (Király et al., 2014) | |
| 14.713601 | 46.170122 | Slovenia | Paper | (Šajna et al., 2017) | |
| 13.935556 | 45.61 | Slovenia | Paper | (Šajna et al., 2017) | |
| 36.117732 | 34.820993 | Syria | Flora | (Mouterde, 1983) | |
| 36.757834 | 35.13179 | Syria | Flora | (Mouterde, 1983) | |
| 10.6346 | 35.8245 | Tunisia | Paper | (Brandes, 2001) | |
| 8.75801 | 36.954432 | Tunisia | Flora | (Pottier-Alapetite, 1979) | |
| 10.333333 | 36.866667 | Tunisia | Flora | (Pottier-Alapetite, 1979) | |
| 10.305191 | 36.818251 | Tunisia | Flora | (Pottier-Alapetite, 1979) | |
| 10.34163 | 36.728658 | Tunisia | Flora | (Pottier-Alapetite, 1979) | |
| 10.805477 | 37.217381 | Tunisia | Flora | (Pottier-Alapetite, 1979) | |
| 10.380833 | 36.135278 | Tunisia | Flora | (Pottier-Alapetite, 1979) | |
| 27.386562 | 37.022841 | Turkey | Citizen science | (iNaturalist, 2020) | gbifID 33941208 |
| 27.388901 | 37.02489 | Turkey | Citizen science | (iNaturalist, 2020) | gbifID 33518883 |
| 27.388967 | 37.025 | Turkey | Citizen science | (iNaturalist, 2020) | gbifID 29658853 |
| 34.916042 | 36.927358 | Turkey | Citizen science | (iNaturalist, 2020) | gbifID 18859850 |
| 34.915999 | 36.927426 | Turkey | Citizen science | (iNaturalist, 2020) | gbifID 18859686 |
| 34.91527 | 36.927186 | Turkey | Citizen science | (iNaturalist, 2020) | gbifID 18845394 |
| 28.302778 | 36.836389 | Turkey | Paper | (Tavsanoglu & Gürkan, 2009) | |
| 31.793053 | 41.451392 | Turkey | Herbarium | (GBIF.org, 2020) | gbifID 2514560493 |
| 31.995326 | 36.543513 | Turkey | Herbarium | (GBIF.org, 2020) | gbifID 1230587109 |
| 27.372747 | 37.948564 | Turkey | Herbarium | (GBIF.org, 2020) | gbifID 665829189 |
| 26.833333 | 39.7 | Turkey | Herbarium | (GBIF.org, 2020) | gbifID 665629405 |
| 36.333333 | 41.25 | Turkey | Herbarium | (GBIF.org, 2020) | gbifID 665620712 |
| 27.543504 | 37.443577 | Turkey | Herbarium | (GBIF.org, 2020) | gbifID 178093917 |
| 58.383333 | 37.95 | Turkmenistan | Herbarium | (GBIF.org, 2020) | gbifID 1140626901 |

### Appendix S2: Defining the historic native range limit

*France.* At the start of the 20th century, the northern range limit of *D. graveolens* on the west side of the continent was located in central France, which represents the most temperate climate within the native range at that time. In France, the species was reported by contemporary botanists as present in the west, south-west, Mediterranean and central parts of the country up to Paris, but absent in the north and east (Bonnier & Layens, 1909; Coste, 1903; Rouy, 1903). To define the range limit, we used the geographical regions as specified by regional landscape features used by Bonnier & Layens (1909). We refined these boundaries using a more detailed account of *D. graveolens*' distribution from Fournier (1946), placing the range limit along the line Lyon-la Loire-Cotentin. We then evaluated all GBIF occurrences dated before 1930; for undated observations, we used the publication year of the reference (botanical books or papers). Starting in Cotentin, the historic native range limit follows the edge of the Loire river basin, extending northwards to include occurrences from the 1800s around Paris (Baillon, 1890), the departments Yonne (Moreau, 1873) and Aube (Briard, 1881), and Côte-d'Or (Genty, 1933, occurrence dated 1880). These references report sporadic observations of *D. graveolens*, but generally describe the species to be rare in this region at the time (Briard, 1881; Brisout de Barneville, 1875; Cosson & Germain de Saint-Pierre, 1861), consistent with the nature of a range edge. From the Loire Basin, the range limit extends a little east towards Lyon, following the river Rhône south into the Mediterranean region. The south-east range limit in France is bounded by elevation ranges >500m as mapped by Bonnier & Layens (1909).

*Italy, the Balkan Peninsula, and Anatolia.* In Italy, *D. graveolens* was most common in the south and center of the country, and was historically absent from the northern regions Aosta Valley, Trentino-South Tyrol, and Friuli-Venezia Giulia (Acta Planatarum, 2020). Here, the historic native range limit follows the northern edge of the River Po Basin. It then continues onto the Balkan Peninsula at the Croatian-Slovenian border. In the Balkan region, we follow *D. graveolens*' distribution as described by Hayek & Markgraf (1931), with two additions: (i) the country of Bosnia-Herzegovina based on the Euro+Med PlantBase (von Raab-Straube, 2021) and a herbarium record from 1907 (GBIF.org, 2020), and (ii) all remaining areas of Greece and Albania not included by Hayek & Markgraf (1931), following the present-day Greek border from Albania to the Thrace region. Recent botanical papers confirm the historical absence of *D. graveolens* from Slovenia (Frajman & Kaligarič, 2009), Hungary (Takács et al., 2016), and Serbia excluding Kosovo (Zlatković & Bogosavljević, 2014). From East Thrace, the range limit continues eastward around the coast of Anatolia following the Flora of Turkey and the East Aegean Islands (Davis, 1975).

*Eastern and southern range limits*. The original native range of *D. graveolens* included Mediterranean north Africa and the Levant, continuing east toward India. To define the southern and eastern range limits, we followed the broad distribution sketch for *D. graveolens* by Brullo

& De Marco (2000) and increased our native range envelope to include additional occurrences reported in West Asia. A historical flora of northwest India (Hooker, 1882) confirms the presence *D. graveolens* there at the time. Reflecting a common bias in the botanical literature, historical records for the east and south of the native range are much scarcer than in the European part of the native range. Fortunately, because the primary aim of our study is to explore the northward expansion, these range edges are much less critical to our analysis.

*Sporadic occurrences north of the original range limit.* We located several accounts prior to 1930 of ephemeral occurrences of *D. graveolens* north of the defined historic native range limit. Where more information from the original data sources could be obtained, these species records were typically described as ephemeral and associated with the wool industry (Scotland: Hayward & Druce, 1919; Netherlands: Kloos, 1940, Belgium: Verloove 2019), salt mines, gravel pits, train stations and other anthropogenic activity known to transport seed (Germany and Switzerland: Wagenitz, 1966). We consider these records to be adventitious sink populations and did not include them within the historic native range limit. In Scotland, we found no recent records for *D. graveolens* that were not associated with the location of the former wool factory (Hayward & Druce, 1919), so we excluded all records in Scotland to be conservative in our niche estimate in the expanded native range (last record in Scotland reported for 1969; BSBI, 2020).

**Appendix S3: Target background point selection**

To set up target background points with a similar spatial bias to our occurrences for *Dittrichia graveolens*, we obtained a total of 214,998 presence records for the congeneric species *Dittrichia viscosa* (GBIF.org, 2021a) and the sister genera *Inula* (201 species; GBIF.org, 2021b) and *Pulicaria* (94 species; GBIF.org, 2021c). Species in both sister genera have distributions collectively covering our study area in the Eurasian Holarctic. Using the same procedure as for *D. graveolens*, we thinned the target records down to a spatial grid with a resolution of 0.5° latitude and longitude, yielding 3468 background points in total across the study area (Fig. S1b).

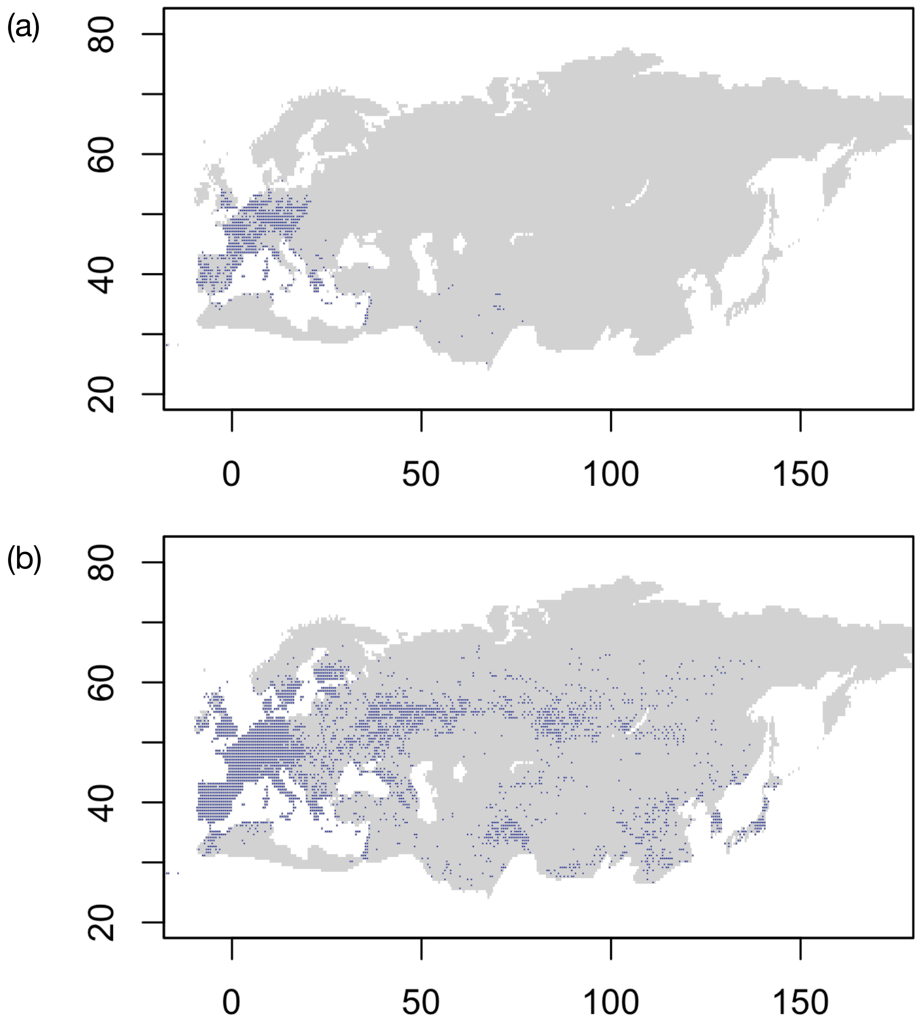


**Figure S1** Presence records for *D. graveolens* (a) and target background points (b).

**Appendix S4: Maxent model projections for the entire Eurasian Holarctic**

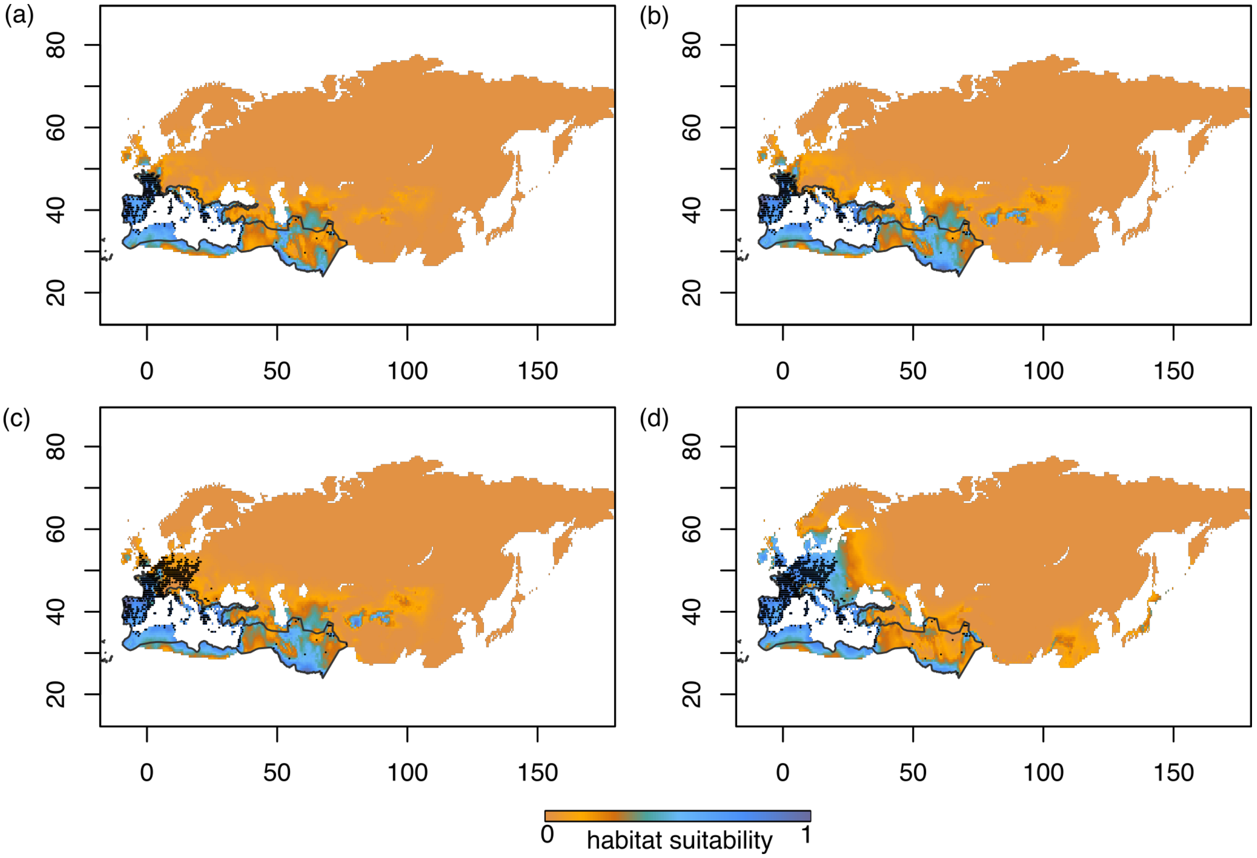


**Figure S2** Maxent habitat suitability results for (a) the Past Model (fit to the past climate and historic native range occurrences) projected onto the past climate (1901-1930); (b) the Past Model projected onto the present climate (1990-2019), indicating expected range expansion with climate change; (c) the same projection with observed occurrences in the present, and (d) the Present Model (fit to the present climate and all occurrences) projected onto the present climate. Historic native range limit represented by the black line, and species occurrence records in the historic (a,b) and expanded (c,d) native range by dots. Axes display decimal degrees longitude (x) and latitude (y).

### Appendix S5: Maxent response functions for each predictor

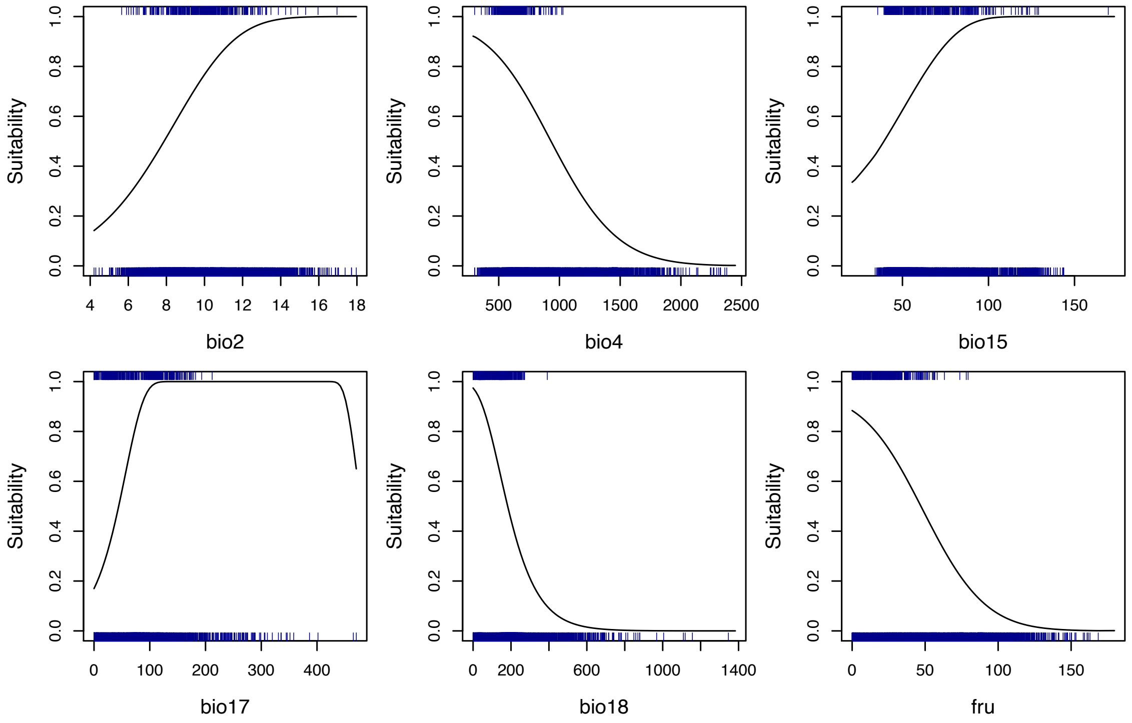


**Figure S3** Response functions for each predictor in the Past Model (Table 1). Blue ticks indicate presences (top) and target background points (bottom). Code adapted from: Smith, A. B. 2020. *Best practices in species distribution modeling: a workshop in R*. http://www.earthskysea.org/.

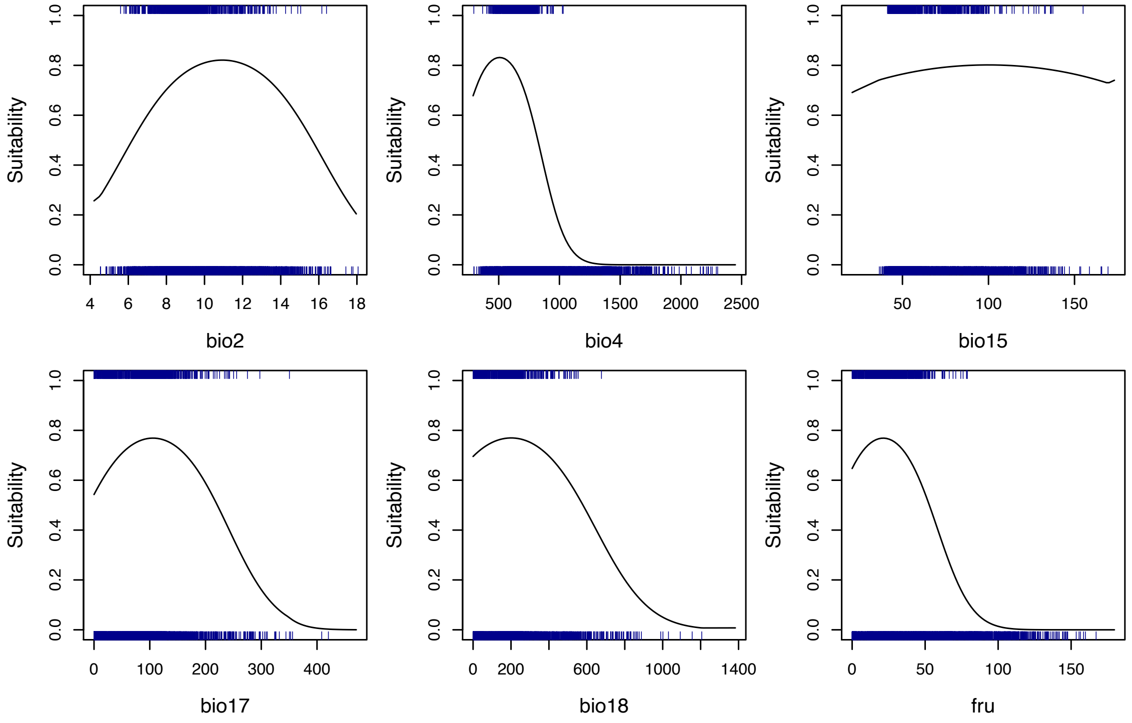


**Figure S4** Response functions for the Present Model, formatted as in Figure S3.
